# The Evaluation of Acute Myeloid Leukaemia (AML) Blood Cell Detection Models Using Different YOLO Approaches

**DOI:** 10.1101/2021.08.04.455113

**Authors:** Kaung Myat Naing, Veerayuth Kittichai, Teerawat Tongloy, Santhad Chuwongin, Siridech Boonsang

**Author notes:** (SB).

## Abstract

This study proposes to evaluate the performance of Acute Myeloid Leukaemia (AML) blast cell detection models in microscopic examination images for faster diagnosis and disease monitoring. One of the popular deep learning algorithms such as You Only Look Once (YOLO) developed for object detection is the successful state-of-the-art algorithms in real-time object detection systems. We employ four versions of the YOLO algorithm: YOLOv3, YOLOv3-Tiny, YOLOv2 and YOLOv2-Tiny for detection of 15-class of AML blood cells in examination images. We also acquired the publicly available dataset from The Cancer Imaging Archive (TCIA), which consists of 18,365 expert-labelled single-cell images. Data augmentation techniques are additionally applied to enhance and balance the training images in the dataset. The overall results indicated that four types of YOLO approach have outstanding performances of more than 92% in precision and sensitivity. In comparison, YOLOv3 has more reliable performance than the other three approaches. Consistently, the AUC values for the four YOLO models are 0.969 (YOLOv3), 0.967 (YOLOv3-Tiny), 0.963 (YOLOv2), and 0.948 (YOLOv2-Tiny). Furthermore, we compare the best model’s performance between approaches that use the entire training dataset without using data augmentation techniques and image division with data augmentation techniques. Remarkably, by using 33.51 percent of the training data in model training, the prediction outcomes from the model that used image partitioning with data augmentation were similar to those obtained using the complete training dataset. This work potentially provides a beneficial digital rapid tool in the screening and evaluation of numerous haematological disorders.

## Introduction

Leukaemia is one type of blood cancer disease related to the rapid production of abnormal white blood cells that generally originate in the bone marrow and human blood [1]. Acute Myeloid Leukaemia (AML) is one subtype of the four-leukaemia blast cell [2] and it can also be classified into distinctive categories by the type of mature and immature white blood cells (blast cells). Unlike any other type of cancer cell, the stages of leukaemia are differently determined based on blood cell counts and the accumulation of leukaemia cells within the organs [3]. Since the blast cell can be propagated and differentiated at a faster pace, the early-stage detection of Leukaemia can save a lot of lives. Inaccurate diagnosis can have an adverse effect directly on the patient for further prescribing the drug regimen as well as increase the cost of the treatment and also could have resulted in the complicated treatment procedure. The identification of the abnormal white blood cells (WBCs) causing the disease may provide important information to physicians in the determination of the appropriate treatments [4]. A microscopic examination method is an early method for categorizing and justifying the cause of the disease. Examination results could be varied depending on sample preparation processes and also the experience and personnel-training of examiners, which lead to inter-and intra-variation of the examining results [5, 6]. Despite the fact that this procedure is carried out by medically qualified examiners who are familiar with the analysis of medical evidence, it has the possibility to be misdiagnosed owing to its subjectivity and the ambiguity of AML blast cell characteristics. It is also challenging to assign standardized images with the time constraint. As a result, the study of blast cell classification is important in early screening of medical diagnosis for further effective planning of the prescription drug regimen.

Several image processing algorithms were formerly applied in mature leukocytes cell image segmentation and classification tasks. Rezatofighi et al. proposed the Gram-Schmidt orthogonalization with snake algorithm to recognize five classes of WBCs and its overall segmentation accuracy is 93% [7]. Discriminating region finding method with three classifiers namely multilayer perceptron (MLP), support vector machine (SVM) and hyperrectangular composite neural networks (HRCNN) was used for recognition of five classes of WBCs and its overall classification rates are 99.11%, 97.55% and 88.89% respectively [8]. Wang et al. proposed a spectral and morphologic method for identification of mature WBCs and the overall accuracy for five classes of WBCs was over 90% [9].

Moreover, Kothari et al. presented the concavity detection and ellipse fitting techniques for the cell boundaries detection and cell counting in digital tissue images sample [10]. The Acute Lymphoblastic Leukaemia (ALL) cell images were identified using this technique by extracting concave points on contour evidence extrication and estimating the contour segment grouping of the leukocytes cells [11]. The concave point detection was also used to separate the overlapped region in acute leukaemia cells [12] and the concave-convex iterative repair algorithm was applied for the segmentation of leukocytes cell images [13]. Although the above techniques have well utilization in accurate definition of cell boundaries and separation of overlapping objects, it is not trivial for the limited resolution of the noisy images.

Through the accelerated adoption of deep learning approaches in real-world applications, various approaches of medical science have emerged in the healthcare sector, using massive amounts of medical data such as electronic health reports, emails, images, sensor data, imaging, and so on [14]. Although the deep learning has gained remarkable success in medical applications, there are still many challenging problems due to the usage of different equipment, the data quality, data availability, and so on. Medical imaging is one of the most important part in medical data and deep learning algorithms are widely applied in medical image analysis [15].

Convolutional Neural Networks (CNN) are neural network architectures that are commonly used to solve image recognition and registration problems. The combination of a CNN and RNN (Recursive Neural Network) model was proposed for the classification of four classes of mature WBCs [16]. The researchers modified four kinds of CNN-RNN models, and the combination of Xception-Long Short-Term Memory (LSTM) has the highest classification accuracy of 90.79%. Cell3Net (deep residual neural network-based leukocyte classifier) was also proposed to recognize 40 categories of WBCs based on its classifications and cytometry [17]. Their results demonstrated the overall accuracy of the test set nearly 76.84%. Jiang et al. also adopted WBCNet to recognize 40 categories of WBCs and average accuracy is 83% on their test dataset [18]. The combination of CNN Fine-Tuning classifier and CNN-feature extraction with SVM classifier based on AlexNet for classifying five classes of mature WBCs was also reported [19]. They presented the inspiring results of average accuracy performance on a test set of both classifiers of 96.38% and 95.12% respectively. Four types of deep learning models namely, AlexNet, ResNet50, DenseNet201, and GoogleNet were used to classify four types of WBCs [20]. Besides, the researcher compared the results by applying Gaussian and median filters separately to the images in the database. The results showed that the applying of Gaussian filter obtained the highest accuracy (83.44%) in the image classification tasks. Acevedo et al. proposed VGG-16 and Inceptionv3 model with SVM classifier to classify 11 classes of leukocytes cell which included mature, immature and platelets cell [21]. The accuracy values for both classes obtained 96% and 95% respectively. Jung et al. also proposed that W-Net, a CNN-based WBC classification model achieved average accuracy at 97% in discriminating of five types of WBC recognition [22].

Moreover, Kutlu et. al. presented the five classes of mature leukocyte cell images detection and classification with the full learning and transfer learning of AlexNet, VGG 16, GoogLeNet, ResNet CNN architecture base on Regional Convolutional Neural Networks (R-CNN) [23]. The result shows that the prediction accuracy of five classes of mature leukocyte cell images achieved more than 94% in transfer learning. Wang et. al. proposed two well-known CNN-based object detection approaches, Single Shot Multibox Detector (SSD) and You Only Look Once Version3 (YOLOv3) for 11 categories of peripheral leukocytes recognition [24]. The two approaches also demonstrated good classification results with a mean accuracy of 90.09% for SSD and 89.36% for YOLOv3. From the above studies, we noticed that the leukocyte cells have more categories depending on their characteristics. The stage identification of the leukocyte cell may assist the prediction of the patient’s health and decide on the treatment process. This fact encouraged us to search for the relatively large dataset which has more stage categories in leukocyte cells by applying a deep learning approach. Matek et. al. published the report on the recognition of blast cell in AML with ResNeXt CNN classification for 15 classes of leukocyte cell images [25]. After five-fold cross-validation, their proposed deep learning networks achieved excellent results in the classes which had more than 400 images in each class. For some classes which have fewer than 100 images, the classification results were still challenging. The balanced sample sizes for each class are critically essential for the CNN model training. Therefore, we propose the four types of the well-known object detection approach, YOLO (You Look Only Once) algorithm to detect the 15-class of WBC cell images. Data augmentation techniques are also employed to increase and balance the training images in the dataset. Furthermore, we compare the two YOLOv3 approaches; one approach uses the entire training dataset and does not use data augmentation strategies, and our proposed methodology is used to determine the gaps in model performances. We also introduce the performance evaluation technique using the ROC curve. This technique can help to examine the performance of our approaches with the quantitative AUC values. The rest of the paper is organized as Section 2 to Section 4. Section 2 presents the dataset arrangement, data augmentation techniques and four types of YOLO approach with their architectures and explanations. Section 3 provides the discussion of classification results and ROC analysis in detail. Finally, we concluded our study in Section 4.

## Materials and methods

### Dataset and labelling

Dataset for this study was received from The Cancer Imaging Archive (TCIA). The peripheral blood smears were collected from 100 patients diagnosed with Acute Myeloid Leukaemia (AML) at Munich University Hospital between 2014 and 2017. The Munich AML Morphology Dataset contains 18,365 single-cell images and a trained examiner experienced in leukocyte cell classified 15 classes for training and evaluation. Each single-cell image has the size of 400 × 400 pixels (corresponding to approximately 29 μm × 29 μm) including background components such as erythrocytes, platelets and cell fragments. The number of images containing in each class is shown in (Fig 1). The full single-cell image dataset and corresponding annotations are publicly available at TCIA [26].

**Fig 1.**
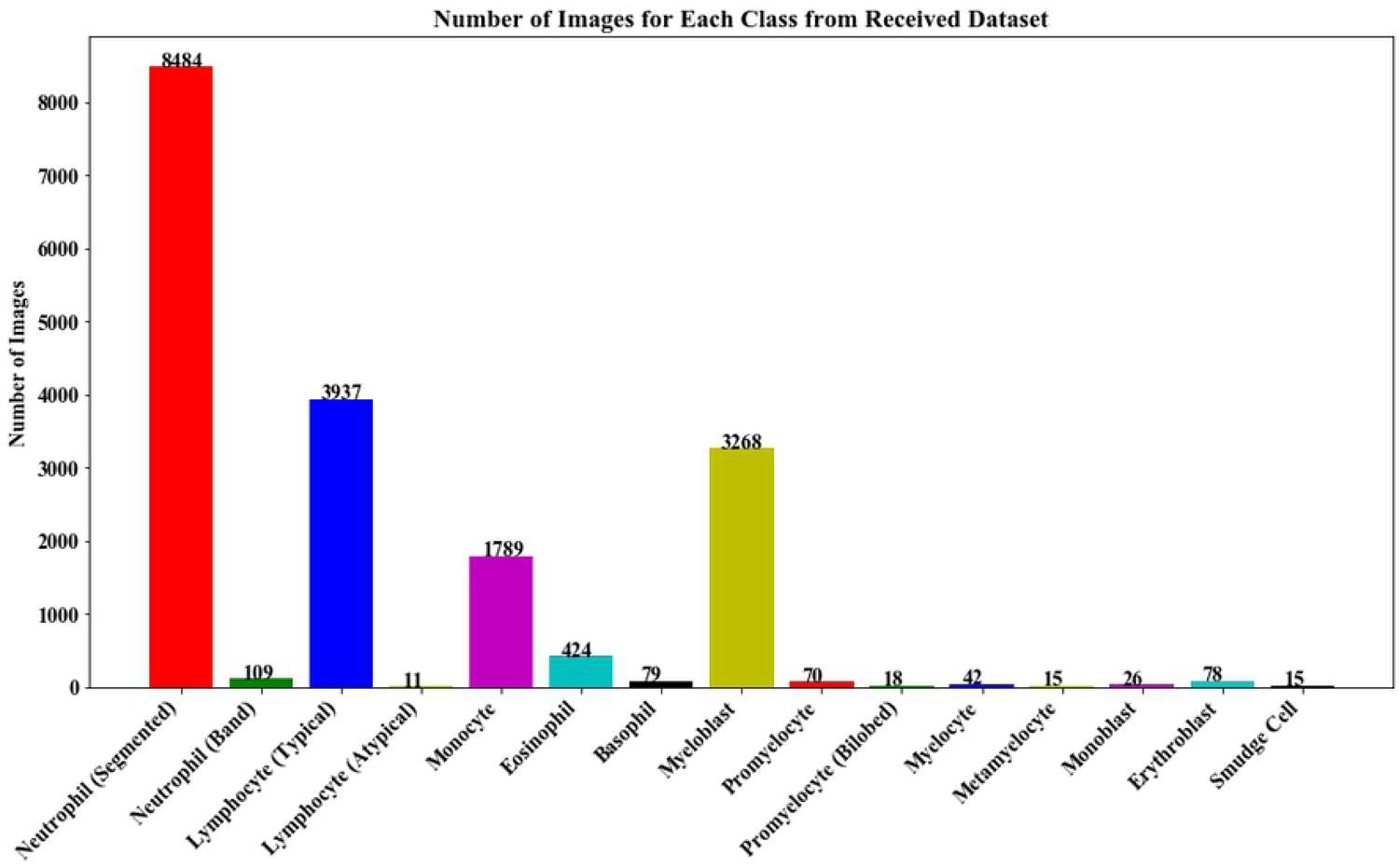
Number of Images for each class from Received Dataset.

According to previous research [27], the most important aspect in machine learning models is the preparation and testing phase. Using different sampling strategies, the researchers investigated the impact of the training and testing processes on machine learning efficiency. The findings revealed that if the dataset is small or perhaps the number of samples in the dataset is small, the test rate can be set between 10% and 20%. In general, the test rate ranges between 20% and 50%. So, we randomly divide the received dataset into a training dataset and testing dataset for each class where the training dataset contains about 80% and the remaining dataset is testing dataset as shown in (Fig 2(a)). To offset the number of images in the dataset, we must minimize certain classes in the testing dataset that have a large number of images. As a result, the three classes of over 3000 images were reduced: Neutrophil (Segmented) was reduced to 1510 images, and the other two classes, Lymphocyte (Typical) and Myeloblast, were reduced to 1000 images each. The number of data analysis for training, testing and unused images is shown in (Fig 2(b)) and the training dataset is reduced from 80% to 27% in entire dataset according to the pie chart.

**Fig 2.**
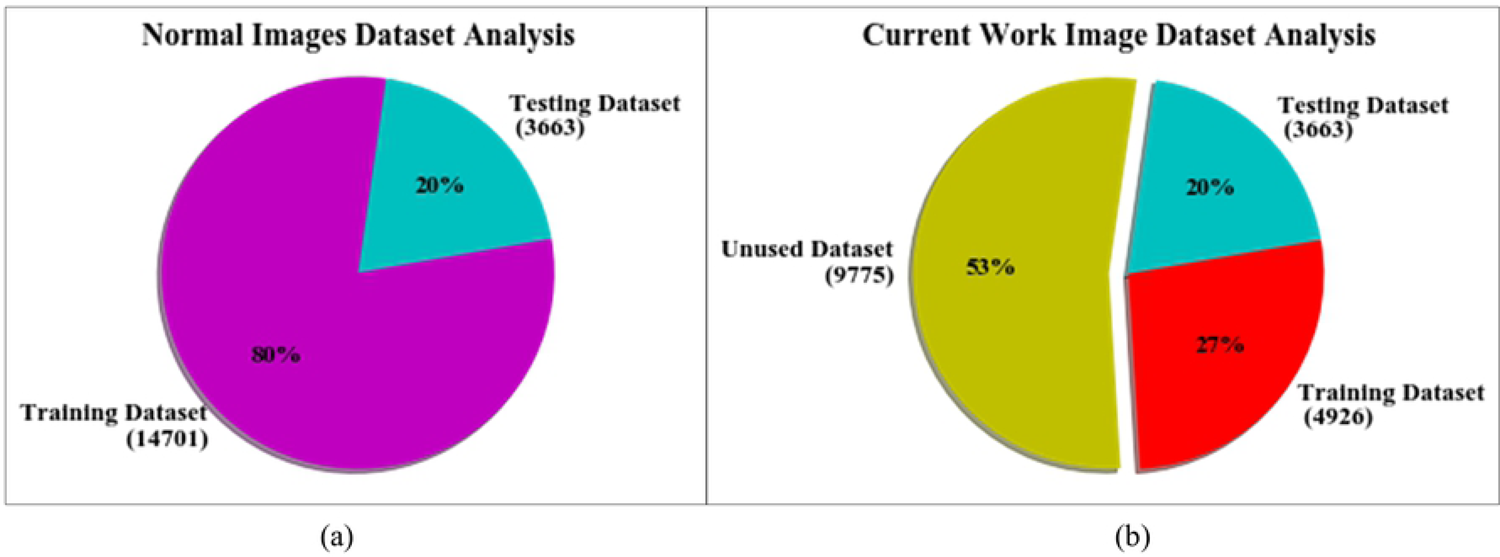
Number of Images Analysis. (a) Number of Images used from received dataset; (b) Number of images used in this research.

Moreover, the detail data utilization of AML dataset is described in Table 1. According to the Table 1, we noticed that the four classes namely Lymphocyte (atypical), Promyelocyte (bilobed), Metamyelocyte, Smudge cell have lower than 20 images and the testing data set has only 2 and 3 images. To evaluate these four classes, the datasets are too small and difficult to get the actual performances.

**Table 1.** Data utilization of acute myeloid leukaemia (AML) blood cells images.

| Classes | Abbreviation | No. of Total Image |  |  | No. of Image Used in this study |  | Usage (%) of Training Data |
| --- | --- | --- | --- | --- | --- | --- | --- |
|  |  | Total | 80% | 20% | Training | Testing |  |
| Neutrophil (segmented) | NGS | 8484 | 6788 | 1696 | 1510 | 1696 | 22.25 |
| Neutrophil (band) | NGB | 109 | 88 | 21 | 88 | 21 | 100.00 |
| Lymphocyte (typical) | LYT | 3937 | 3150 | 787 | 1000 | 787 | 31.75 |
| Lymphocyte (atypical) | LYA | 11 | 9 | 2 | 9 | 2 | 100.00 |
| Monocyte | MON | 1789 | 1432 | 357 | 700 | 357 | 48.88 |
| Eosinophil | EOS | 424 | 340 | 84 | 340 | 84 | 100.00 |
| Basophil | BAS | 79 | 64 | 15 | 64 | 15 | 100.00 |
| Myeloblast | MYO | 3268 | 2615 | 653 | 1000 | 653 | 38.24 |
| Promyelocyte | PMO | 70 | 57 | 13 | 57 | 13 | 100.00 |
| Promyelocyte (bilobed) | PMB | 18 | 15 | 3 | 15 | 3 | 100.00 |
| Myelocyte | MYB | 42 | 34 | 8 | 34 | 8 | 100.00 |
| Metamyelocyte | MMZ | 15 | 12 | 3 | 12 | 3 | 100.00 |
| Monoblast | MOB | 26 | 21 | 5 | 21 | 5 | 100.00 |
| Erythroblast | EBO | 78 | 64 | 14 | 64 | 14 | 100.00 |
| Smudge cell | KSC | 15 | 13 | 2 | 12 | 2 | 92.31 |
| Total |  | 18365 | 14701 | 3663 | 4926 | 3663 | 33.51 |

The method of manually labelling the regions of an object in an image and generating text-based definitions of those regions for object recognition is known as image labelling. Well-trained individuals completed the image labelling description.

The main objective of the image labelling is to allow the users to highlight or specify the pixel area of the objects in an image. The CiRA CORE platform [28] was used as a tool for images labelling before applying data augmentation techniques.

### Data augmentation

Data augmentation is the most commonly-used techniques which were extensively utilized to increase the sample size of a dataset for the model training process. Many researchers used numerous image transformation techniques, such as position augmentation and color augmentation, to obtain various types of original images. Since deep learning models were trained with both the original and various augmentation images, they have more generalization capabilities. As a survey result on image data augmentation for deep learning, various data augmentation techniques have been developed to lessen overfitting of the learning model by providing better generalization [29]. Therefore, we used the following four data augmentation techniques:

- Rotation: It is one kind of position augmentation technique. It was done by giving the rotating effect to the image with the degree value specified from −180° to 180°. In this study, the rotating effect was varied at every 45° to obtain various images as shown in (Fig 3(b)).
- Contrast: It is one kind of augmentation technique in image processing and it can be adjusted between the darkest and brightest image areas. The contrast of images was changed by multiplying all pixel values with 0.4, 0.6, 0.8 and 1.0 as shown in (Fig 3(c)).
- Noise: Noise can reduce the accuracy of neural networks when testing on real-world data. Image noise injection is an important augmentation step that allows the model to discriminate original image from noise in an image. Gaussian noise distribution used are three values of standard deviation (σ = 0, 10, 20) as shown in (Fig 3(d)).
- Blur: Blurring is a very popular technique in image processing. It can be defined as the degree of separation between sharpening and blur images. Gaussian blur filter used will result in a blurrier image. Blurring images for data augmentation could lead to higher resistance to motion blur during testing. The higher the standard deviation (sigma) value, the more will be the blurring effect. Gaussian filter with a standard deviation of 9 was used to blur the images as shown in (Fig 3(e)).

**Fig 3.**
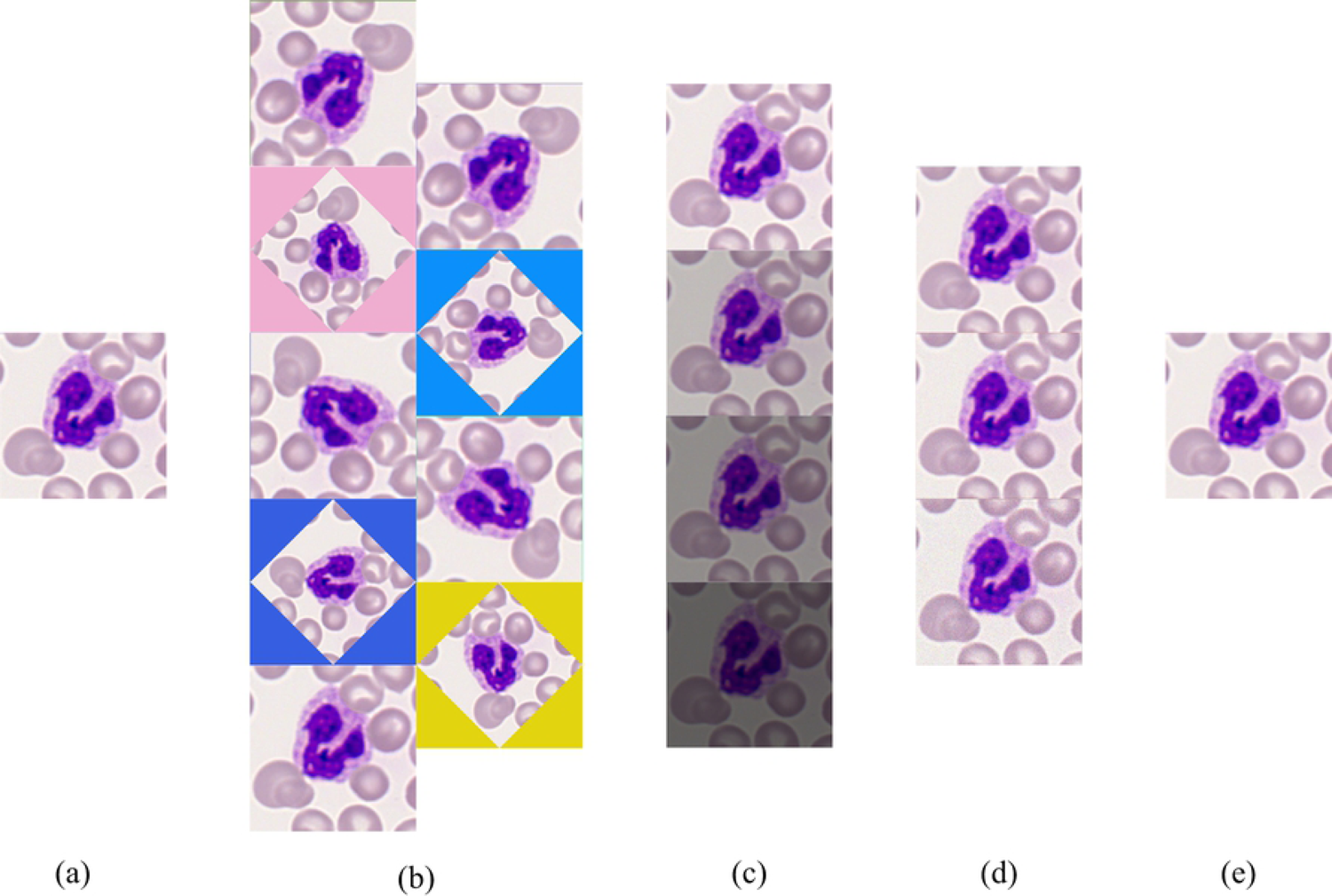
Four Types of Data Augmentation Techniques. (a) Original Image; (b) Image Rotation; (c) Image Contrasting; (d) Noise Injection to Image; (e) Image Blurring.

After completing the four image processing tasks mentioned above, a single image can be augmented into up to 108 images of 608×608 pixel resolution. The testing dataset with the representation of 608×608 pixels was used to prepare the training dataset for the four versions of the YOLO algorithm.

To assess the improvements in our methodology that used data utilization and data augmentation techniques, we used a performance comparison with an approach that used the whole testing dataset without using data augmentation techniques. This analysis is based on the best YOLO approach in the average results analyses of four different YOLO approaches. Image labelling and image pixel selection (608×608) are the same procedures employed in this process’s training as mentioned above.

### Detection and classification of acute myeloid leukaemia (AML) blood cell images based on YOLO approach

The four models of YOLO approach (YOLOv3, YOLOv3-Tiny, YOLOv2 and YOLOv2-Tiny) which used for classification of AML Blood Cell are described as the following subsections.

### Detection and classification of acute myeloid leukaemia (AML) blood cell images based on YOLOv3

In this study, the proposed model for AML Blood Cell classification task is based on YOLOv3 convolutional neural network as shown in (Fig 4).

**Fig 4.**
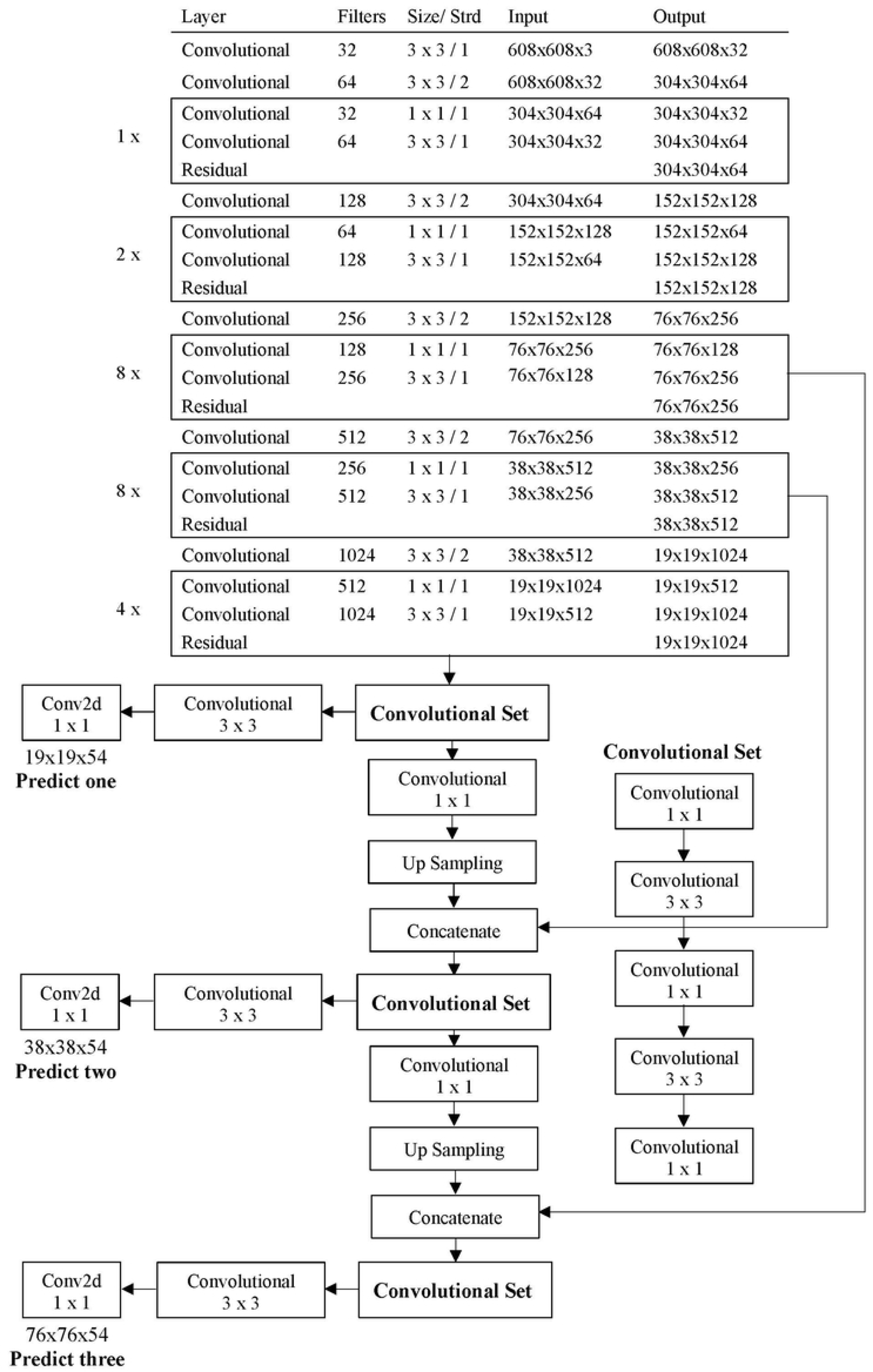
The proposed model of YOLOv3.

YOLO algorithm, one of the most popular state-of-the-art object detectors in real-time object detection, was introduced in 2015. Although the first version of YOLO (YOLOv1) was faster in processing time, it has a lower accuracy than other models at that time [30]. Nevertheless, it has much improved in object detection with the later modified version. The third version of its (YOLOv3) model is one of the preeminent object detectors because it is faster and more accurate than other state-of-the-art models [31]. The YOLOv3 model differs from the previous YOLO versions in its architecture composed of: (i) the Darknet-53 backbone, which is used for feature extraction because it has fewer floating-point operations and more speed than ResNet-101 or ResNet-152, (ii) ResNet short cut connections, which are also added to avoid the disappearance of gradients and (iii) several convolutional layers, which are added in base feature extractor for predicting three different bounding boxes using a similar concept to feature pyramid networks (FPN). Meanwhile, YOLOv3 predicts bounding boxes using the sum of squared error loss and the logistic regression function. Furthermore, for multilabel class predictions, discrete logistic classifiers are used instead of Softmax, and binary cross-entropy loss functions are used instead of multiclass loss functions. The YOLOv3 architecture also uses k-means clustering to evaluate the bounding box priors.

As illustrated in (Fig 4), YOLOv3 finally predicts three-branch outputs in the feature map for each cell image. We selected 608×608 for the input image sizes, and the output has three different types of attribute maps: 19×19 for a large object, 38×38 for a medium object, and 76×76 for a small object. In the training stage of YOLOv3 model using the training dataset, we used the following network parameter: the batch size of 64, maximum batches of 500200, and subdivisions of 16, a momentum of 0.9, a weight decay of 0.0005 and activation function of leaky ReLU (Rectified Linear Unit) for convolutional layer and linear for shortcut (Residual) layer. By default, we adopt the multistep learning rate policy with a base learning rate of 0.001, a step value of [40000, 45000] and the learning rate scales of [0.1, 0.1]. Finally, the output layer predicts both class probabilities and the coordinates of the bounding box using threshold level and NMS (Non-Maximum Suppression) technique.

### Detection and classification of acute myeloid leukaemia (AML) blood cell images based on YOLOv2

To understand more about the improvement of YOLOv3’s performance, we employed YOLOv2 for comparison in this study. YOLOv2 and YOLO 9000 was proposed by J. Redmon and A. Farhadi in 2016 [32]. YOLOv2 is also state-of-the-art algorithm and faster than other detection systems across a variety of detection datasets at that time. The proposed YOLOv2 is used Darknet-19 architecture which composed of 19 convolutional layers and 5 max-pooling layers because it has lower processing requirement than other architectures such as VGG-16 and customized GoogleNet. The proposed architecture of YOLOv2 is shown in (Fig 5).

**Fig 5.**
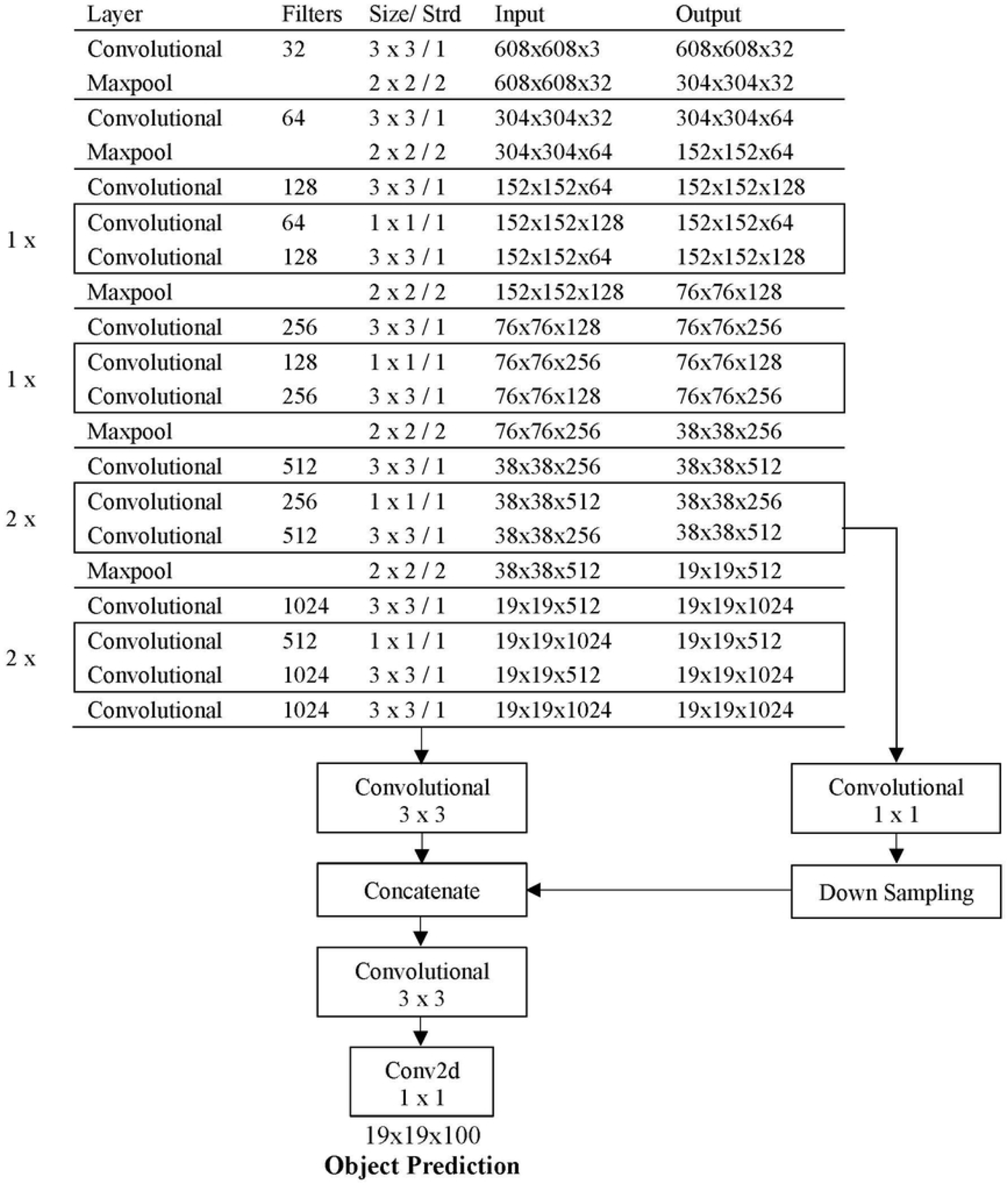
The proposed model of YOLOv2.

In the training of YOLOv2, we used the same network parameters as YOLOv3 except for Softmax classifiers. Furthermore, YOLOv2 gives one output feature map of image size 19 × 19 for object prediction with 5 anchor boxes. The size of these five anchor boxes can be varied depending on training dataset. A list of 10 numbers for these five anchor boxes in this study was: 0.57273, 0.677385, 1.87446, 2.06253, 3.33843, 5.47434, 7.88282, 3.52778, 9.77052, 9.16828. Besides, we selected the size of 608×608 for input image as showed in (Fig 5).

### Detection and classification of acute myeloid leukaemia (AML) blood cell images based on YOLOv3-tiny

YOLOv3-tiny, a simplified version of YOLOv3, is widely used because it runs faster and takes less memory. An advantage of YOLOv3-tiny is that it can test on CPU instead of GPU. The training network parameters are the same as YOLOv3 and the proposed model is showed in (Fig 6). Unlike YOLOv3, three clusters are selected, and the target is distributed into two scales by using k-means clustering. The size of these three clusters is arranged from small to large as follows: 10,14, 23,27, 37,58, 81,82, 135,169, 344,139. The size of 416 × 416 images is selected as input and YOLOv3-tiny finally has two branch outputs for prediction with the size of feature maps 13 × 13 and 26 × 26, respectively.

**Fig 6.**
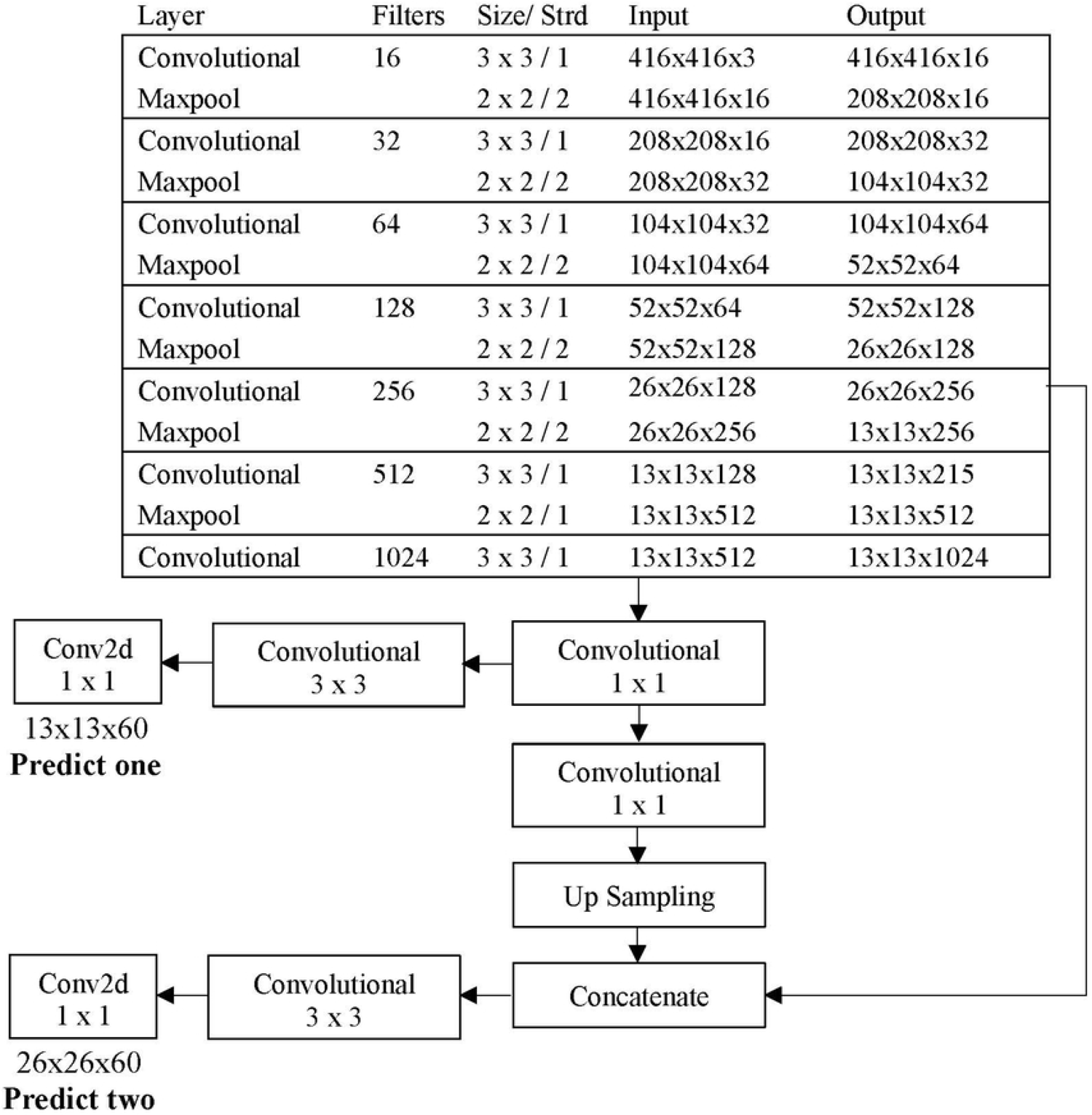
The proposed model of YOLOv3-Tiny.

### Detection and classification of acute myeloid leukaemia (AML) blood cell images based on YOLOv2-tiny

YOLOv2-tiny is a smaller version of the second version of YOLO. It is also a real-time object detection model and is used to compare with YOLOv3-tiny. Its structure is slightly different from YOLOv2 and YOLOv3-tiny. It composed of 9 convolution layers and 6 max-pooling layers and predicted one output feature map as shown in (Fig 7). The training network parameters are the same as the above three models such as batch size, maximum batches, subdivision, momentum, weight decay, learning rate, step values and learning rate scales. The values of five anchor boxes are the same as YOLOv2 and the selection of 416 × 416 image size for input is the same as YOLOv3-tiny. Therefore, the model provides one output feature map with a size of 13 × 13 for object prediction.

**Fig 7.**
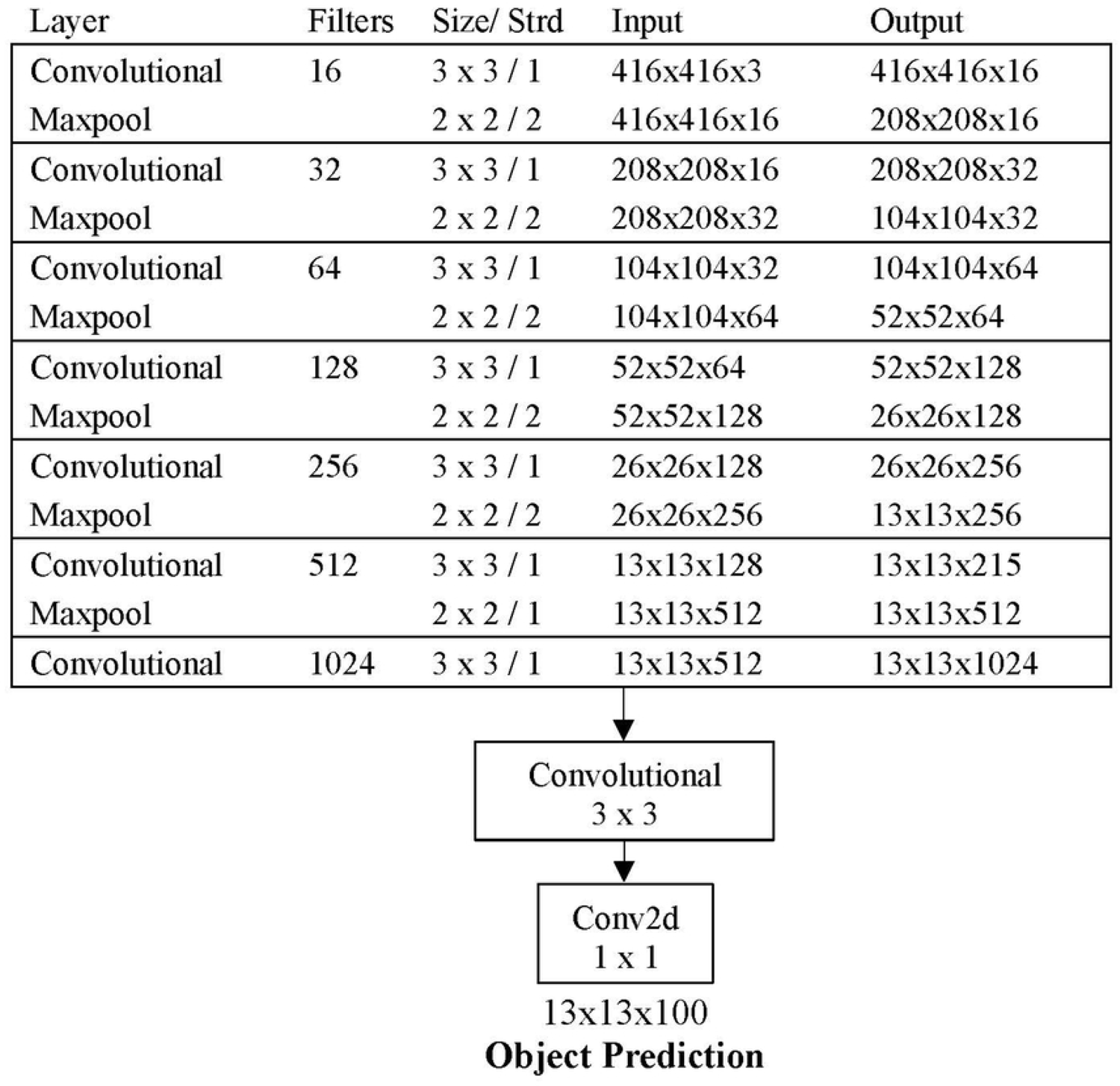
The proposed model of YOLOv2-Tiny.

## Statistical analysis

The performance of the four models is evaluated by using a testing dataset which containing 15 classes of human blood cell images. When we tested the performance for each class, we will get the prediction results with a confidence level. We counted the number of prediction results for each class and summarized together into a 15×15 confusion matrix table. This is because we used 15 classes of blood cell images. The four statistical parameters namely precision, sensitivity, specificity, and accuracy are considered as performance metrics. Besides, the four types of conditions such as True Positive (TP), True Negative (TN), False Positive (FP) and False Negative (FN) are used to evaluate the performance of proposed models [33]. The definition of these conditions in this research are as follows:

- True Class: A class that we require to test the class at that time.
- False Class: The classes that we do not require to test the class at that time.
- True Positive (TP): The model classifies the number of blood cell image correctly when testing the true class.
- False Positive (FP): The model classifies the number of blood cell image correctly when testing the false class.
- False Negative (FN): The model classifies the number of blood cell image incorrectly when testing the true class.
- True Negative (TN): The model classifies the number of blood cell image incorrectly when testing the false class.

After receiving all of these four values for each class, the four-performance metrics as described above are computed using Equations (1), (2), (3), and (4).

Precision is defined as the proportion of true class with a positive test which has true. The precision is calculated by the following formula:

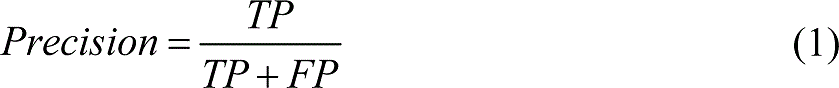

Sensitivity is defined as the proportion of true class which have a positive test. The sensitivity is calculated by the following formula:

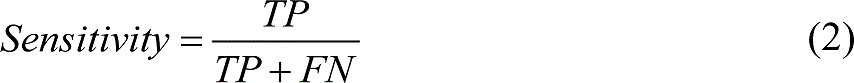

Specificity is defined as the proportion of false class which have a negative test. The specificity is calculated by the following formula:

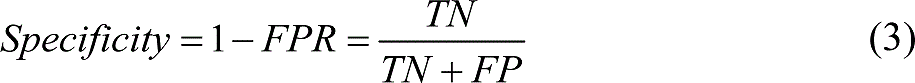

Accuracy is defined as the proportion of both true class and false class which have correctly classified over the entire testing dataset. The accuracy is calculated as follows:

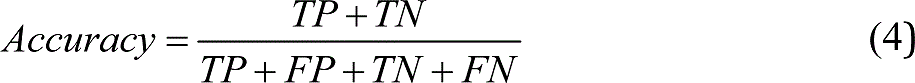

Besides, we can visualize the model performances and evaluate the classifiers with the well-known technique, the ROC (receiver operating characteristic) curve [34]. ROC curve is a two-dimensional graph in which true positive rate (TPR) and false positive rate (FPR) value are plotted on Y-axis and X-axis, respectively. TPR and FPR values can receive from the confusion matrix table to form a ROC curve. TPR value is the same with sensitivity and FPR value can be calculated as follows:

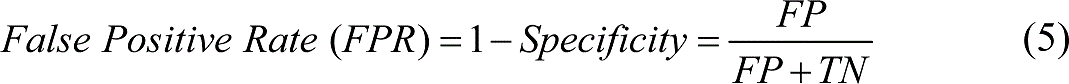

Since the ROC is constructed by changing the threshold level on the prediction score, each changing threshold generates one point in the ROC curve [35]. In this study, we constructed the ROC curve at the threshold level with every 5% increment because our testing data set has over 3000 images. The above description of TP, FP, TN and FN are simply used in the binary class decision. However, we use the multi-class classification with confusion matrix of 15 × 15 for blood cell image analysis. The confusion matrix for n × n is constructed as shown in (Fig 8).

**Fig 8.**
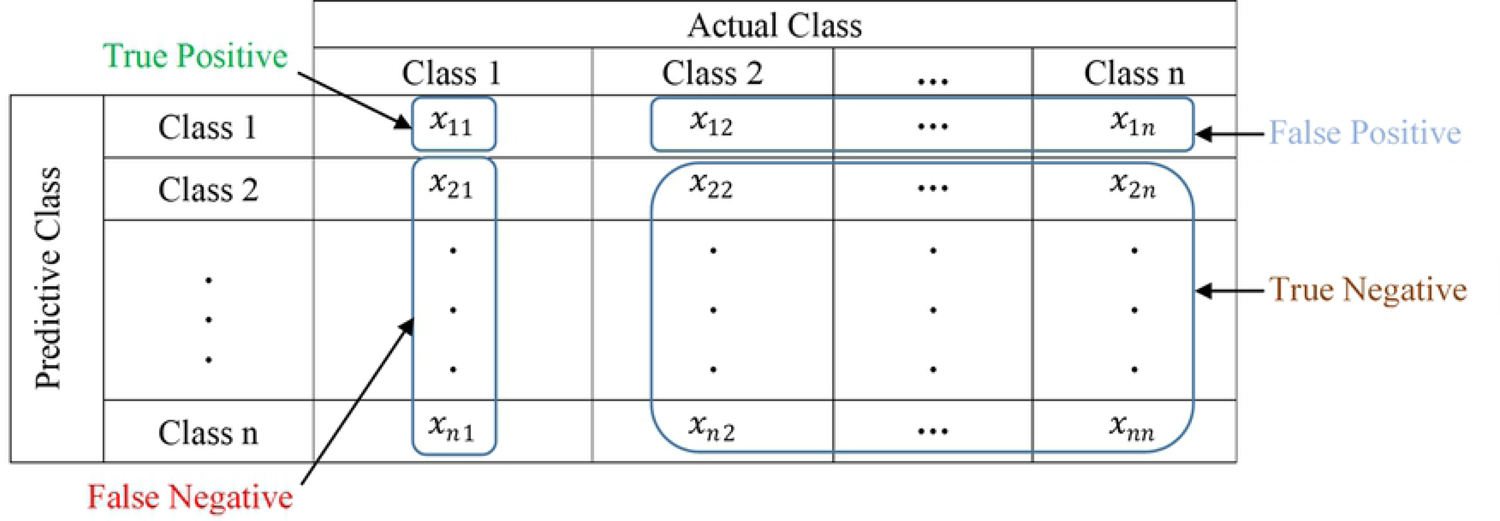
Confusion matrix for *n*-class; where *x* stands for the number of AML cell count for the corresponding conditions and *n* will be 15 classes for this study. For example, the four types of conditions; True Positive (TP), False Negative (FP), True Negative (TN), False Negative (FN) are also described when testing the 1st class of multi-class classification.

In the multi-class classification, the four types of conditions could be computed by using a one-versus-rest approach. Then, the performance calculations can be achieved with two operations, namely micro-averaging and macro-averaging. Because of the imbalance in the class dataset, we used micro-averaging in this study [36]. To plot the ROC curve, the required TPR and FPR are computed as follows:

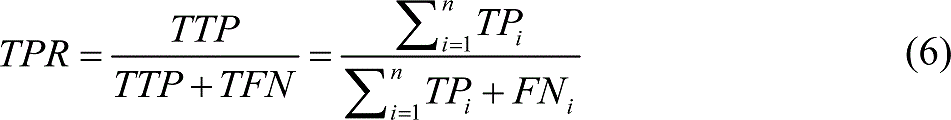

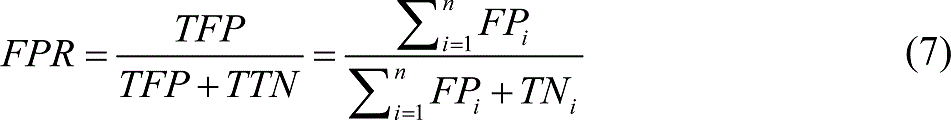

where *i* stands for each testing class and n will be 15 classes for this study. If *i* = 1, the 1^st^ class is considered as positive and the rest of 14 classes as a negative class. In the formulas, *TP_i_*, *FP_i_*, *TN_i_*, *FN_i_* are associated with each testing class *i* and *TTP*, *TFP*, *TTN*, *TFN* are the total of each condition for all classes.

Moreover, we performed the calculation of the area under the curves, or AUC (also known as an AUCROC curve) by the integration of the area under the ROC curve. These AUC values are key measure of the usefulness of the proposed model whereas the greater the curve area means more useful the model in testing.

Since the sensitivity is the same with TPR values, the other three performances metrics on all classes are computed as follows:

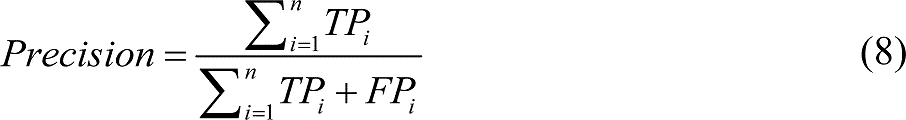

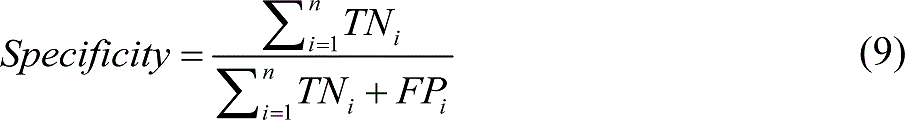

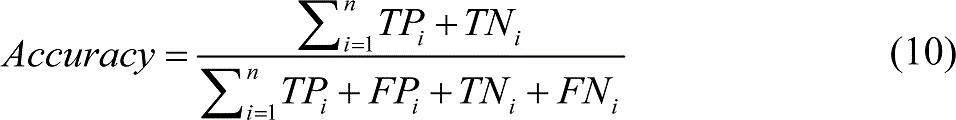

## Results and Discussion

Since YOLO algorithms required a large amount of data, we needed to use a machine that has very high computing power. All of the four algorithms were configured on the operation system of the Ubuntu 16.04 LTS (64-bit) and the hardware environment as follows: Processor: Intel® Core i5-8400, CPU @ 2.8GHz*6, Memory: 31.3 GiB, and Graphics: GeForce GTX 1070 Ti. In this study, the training process took a maximum of 4 days for computing time.

### Performance comparison of classification results on the four algorithms

In this section, we describe the performance comparison for the four YOLO algorithms with the testing dataset. We evaluated the prediction output scores for each class by using the corresponding trained models. The parameter selection of threshold level: 0.5 (50%) and NMS value: 0.2 are used as default to attain the prediction scores. Since there is a 0.5 classification threshold, the models can predict the classes if the probability is greater than 0.5. Besides, we utilized the NMS technique to remove duplication because the model can give duplicate detections output for the same output [37]. After that, we used the testing dataset as described in Table 1 which has 15 classes of blood cell images with a total number of 3,663 single-cell images to achieve the class predictions. We used the actual class label and the predicted class score to produce the confusion matrix table. To complete the confusion matrix, we needed to count the predicted class for each image testing and filled the received counted number on the prediction class of the matrix. The four types of a confusion matrix table for the four YOLO algorithms are shown in (Fig 9).

**Fig 9.**
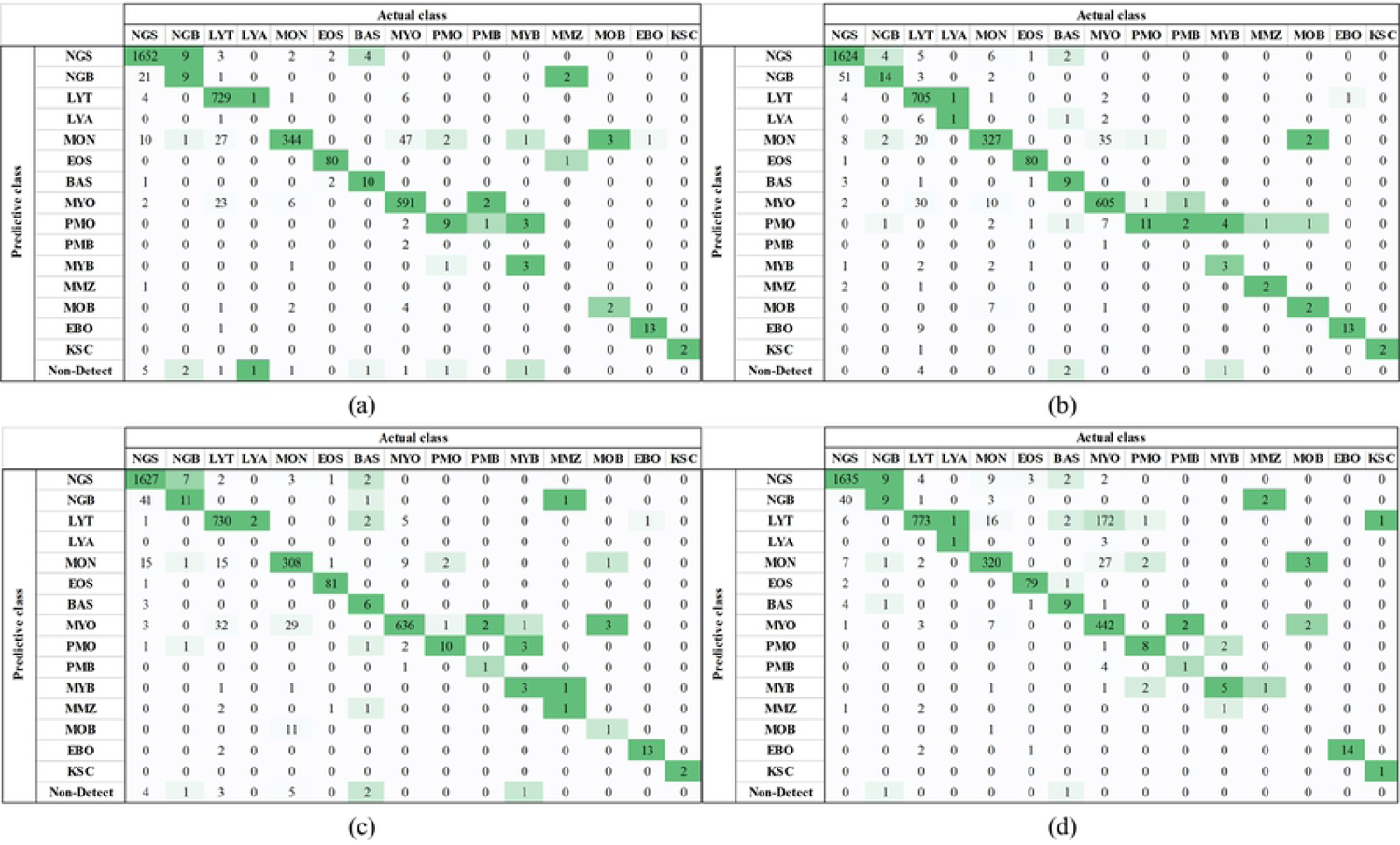
Confusion Matrix for Four YOLO approaches. (a) Confusion Matrix for YOLOv3, (b) Confusion Matrix for YOLOv2, (c) Confusion Matrix for YOLOv3-Tiny, (d) Confusion Matrix for YOLOv2-Tiny. Unlike the general confusion matrix, the non-detect condition is considered in predictive class when the model cannot detect any cell class in AML blood cell image while testing the true class.

In the confusion matrix, we found that the YOLOv2 and YOLOv3-Tiny can correctly predict more classes than the other two classes. These two models can predict 14 classes, the YOLOv2-Tiny can predict 13 classes and YOLOv3 can predict 12 classes even though some class has a small dataset. As mentioned in the Dataset and Labelling section, the four models were difficult to correctly perform in the small testing dataset. Nevertheless, the four models accurately predicted the class namely Smudge cell because it has a distinctive character than other classes. The typical results from the detection and classification of each single-cell images are shown in (Fig 10). Although YOLOv3 predicted fewer classes than the other algorithm, it has more number of prediction images than the other three models. For the total number of image prediction, the four models correctly predicted over 3663 single-cell images as follows: YOLOv3 predict 3444 images, YOLOv3-Tiny predict 3430 images, YOLOv2 predict 3998 images, and YOLOv2-Tiny predict 3297 images, respectively.

**Fig 10.**
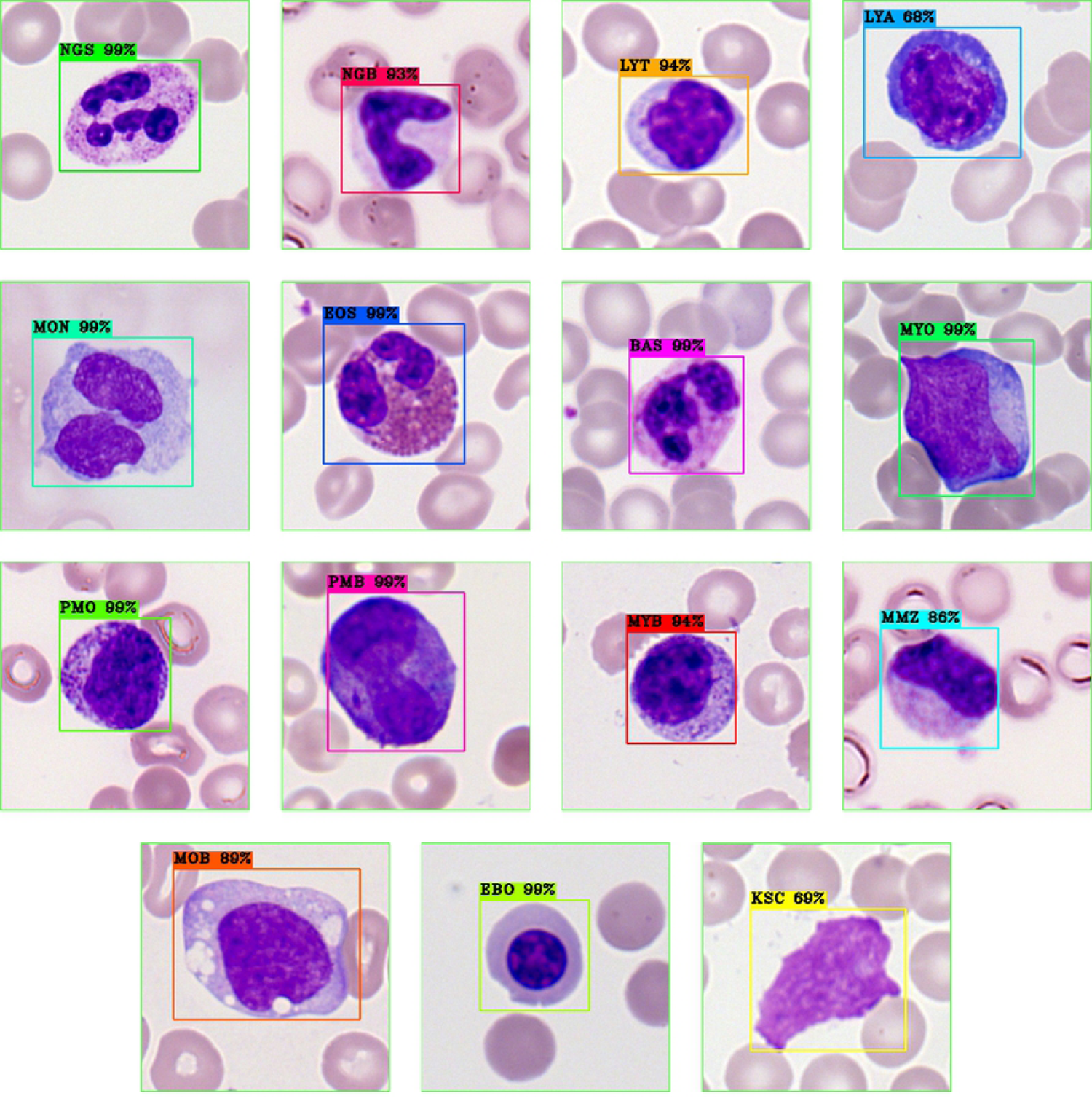
The classification and localization of Acute Myeloid Leukaemia (AML) Blood Cell images with the threshold level of 0.5 and NMS of 0.2.

Additionally, we compare the precision values and sensitivity values to evaluate the quality of class-wise prediction by using one-versus-rest approach as shown in (Fig 11 and Fig 12). At this point, we used the results of the previous work [25] to make the comparison which used the ResNeXt CNN approach for training and 5-fold cross-validation for testing. Consequently, they performed five different times for training and testing. Although they presented their interval values in precision and sensitivity, we only used the average results for comparison.

**Fig 11.**
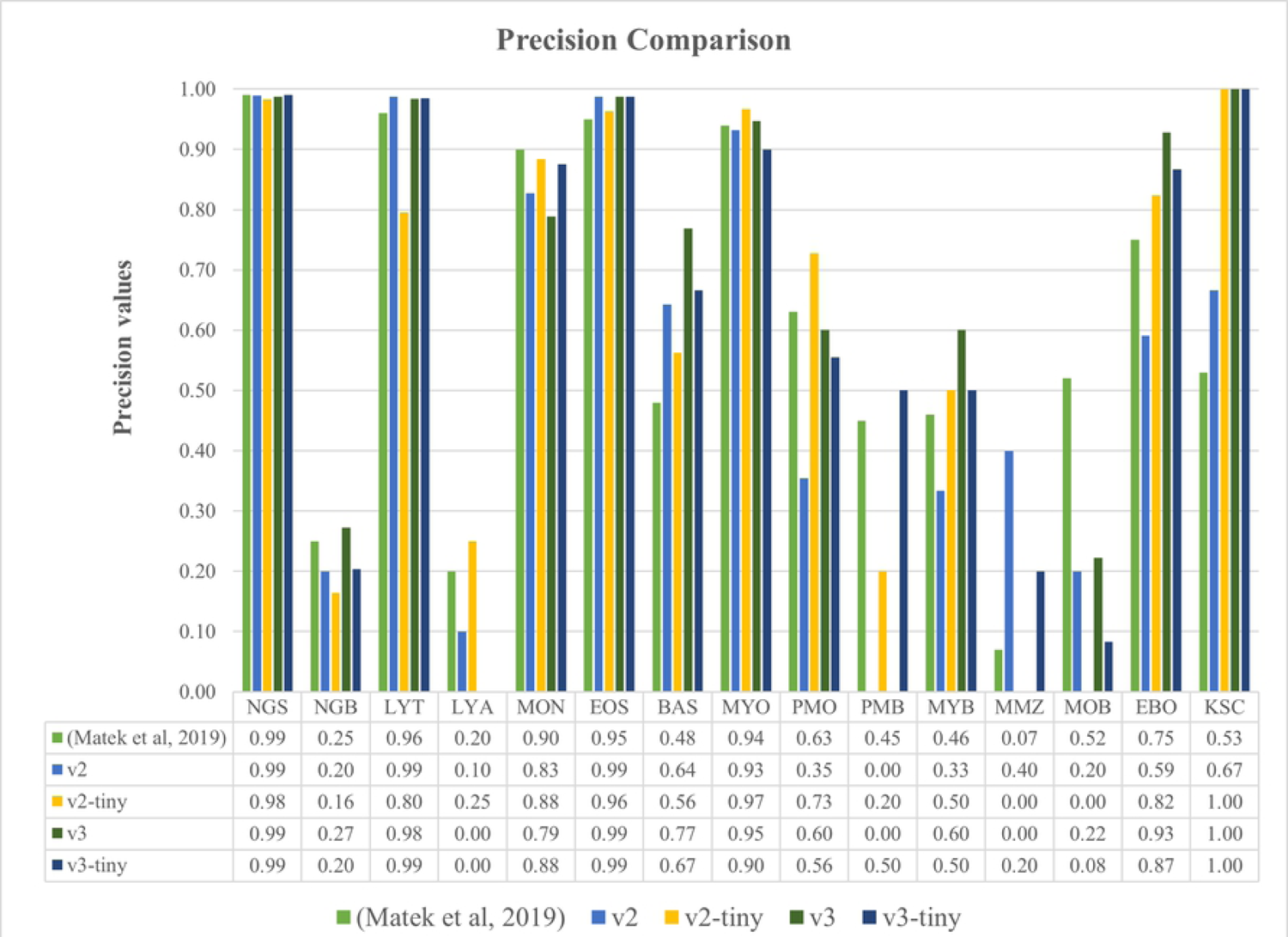
The Comparison of Precision Values for the four types of YOLO and the previous work.

**Fig 12.**
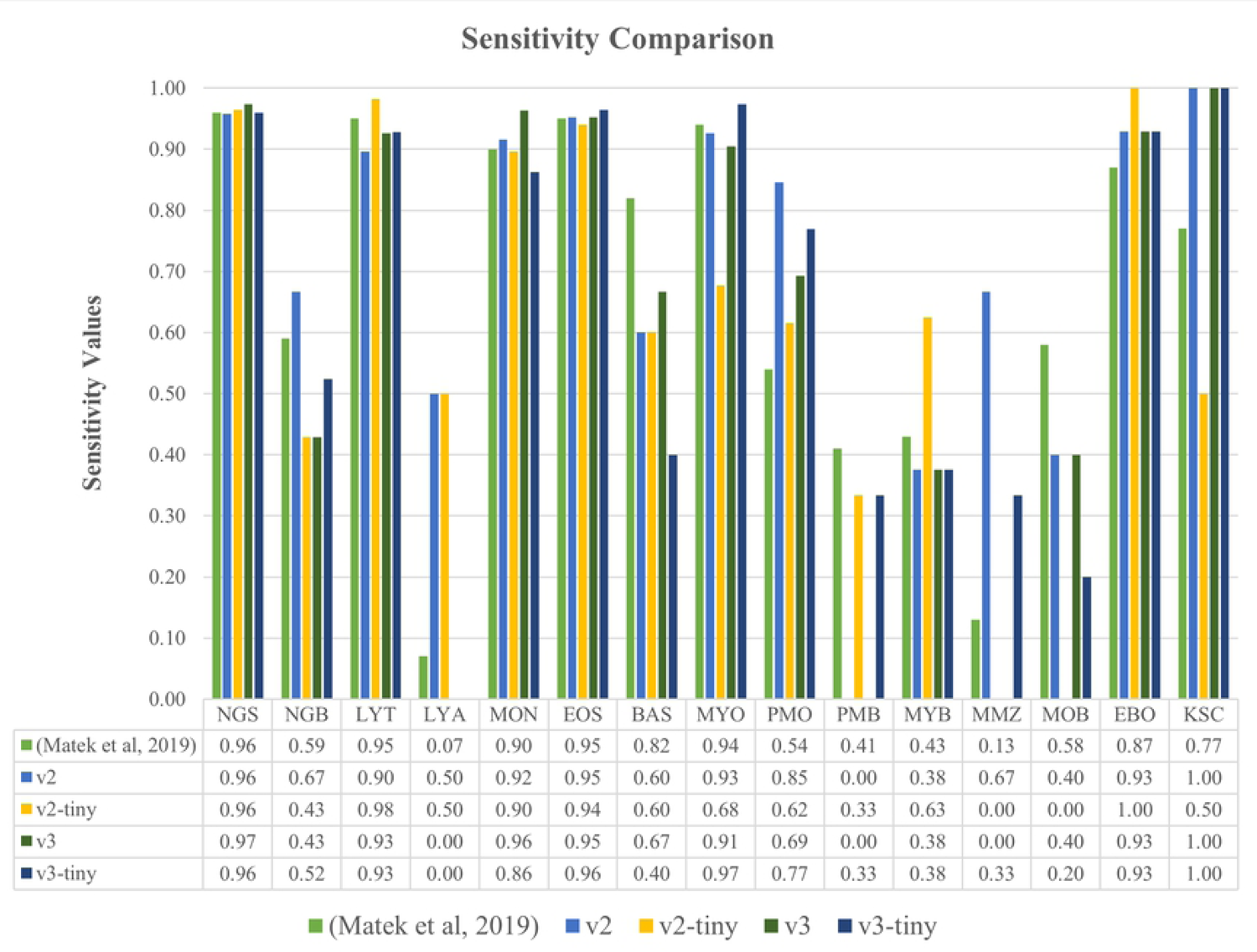
The Comparison of Sensitivity Values for the four types of YOLO and the previous work.

(Fig 11) presents the precision results for the previous work and for four types of YOLO with the bar graph. We found that the three large dataset classes have at least 90% in precision for YOLOv3 and YOLOv3-Tiny. In general, the small dataset classes have low precision values (some classes have 0) except for the Smudge (KSC) cell for all of the models. Because the characteristic of KSC cell is significantly different features without a specific biological pattern from the other classes as shown in Figure (Fig 12) displays the result of sensitivity for all of the models. The best sensitivity scores were obtained at least 90% in the three classes as described in the precision result for YOLOv3 and YOLOv3-Tiny. The two significant characterized classes namely EOS, KSC have relatively high sensitivity score of at least 95%. To summarize the above facts, the large training datasets still provided the capability of YOLO in the classification of blood cell images. Besides, the performance of multi-classification dropped due to the similar characteristic of blood cells for a small dataset. 10. Moreover, Eosinophil (EOS) cell also has a high precision (at least 95%) for all models because of its unique characteristic.

The overall results of four-performance metrics for four types of YOLO models are shown in (Fig 13). YOLOv3 has 94% in overall precision and sensitivity and 99% in overall specificity and accuracy which is a better performance than the other three models. Therefore, we can conclude that the YOLOv3 model is suitable for high-performance GPU whereas YOLOv3-Tiny is compatible with low memory and CPU devices.

**Fig 13.**
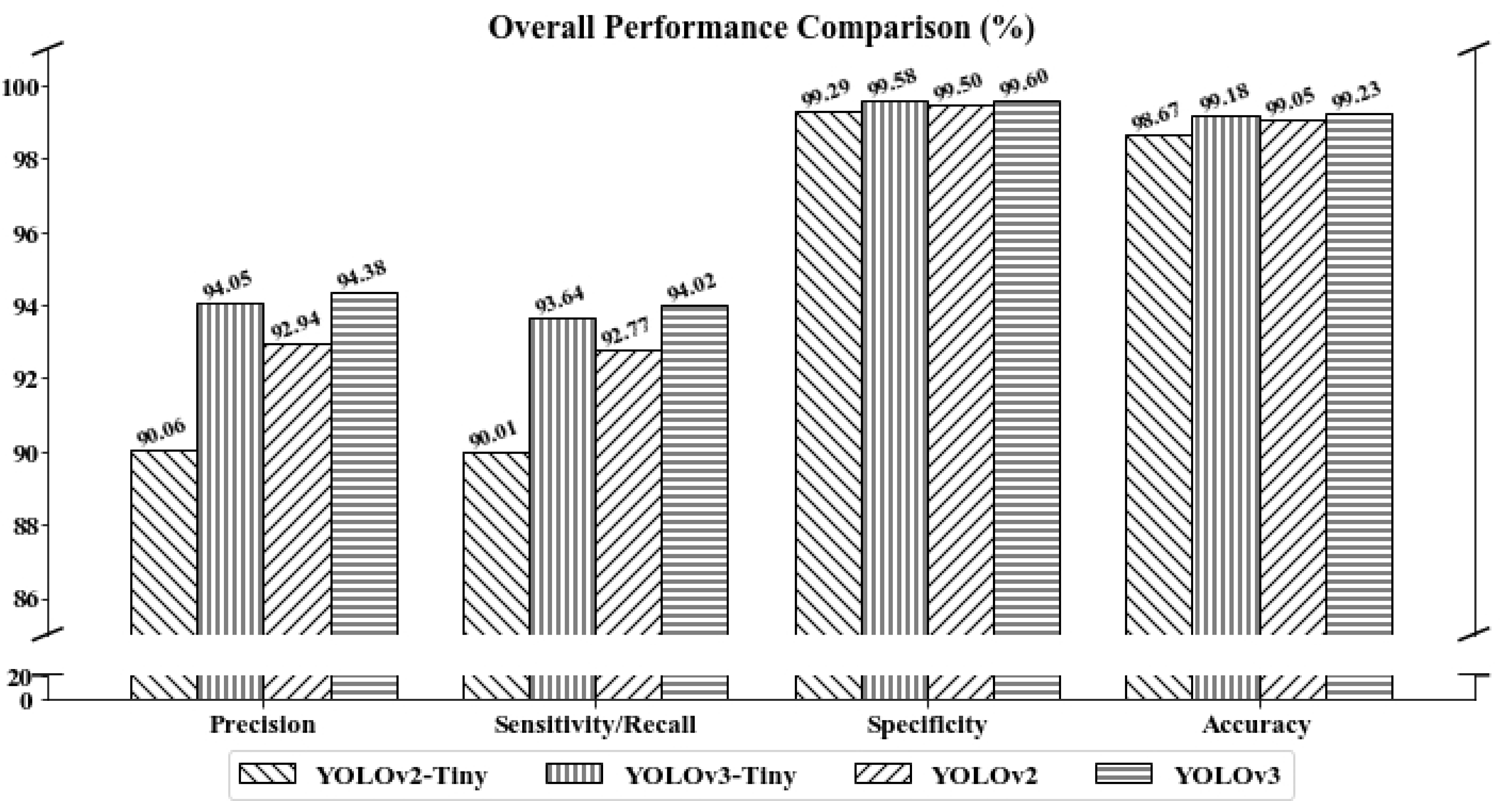
The overall score for the four-performance metrics namely precision, sensitivity, specificity, and accuracy with the threshold level of 0.5 and NMS of 0.2. The model performances are calculated by using micro-averaging.

### The performance comparison between YOLOv3 (using entire training dataset and without applying data augmentation techniques) and YOLOv3 (our approach)

Since the YOLOv3 model, trained with the data partition strategy (our approach), has the best results as defined in the previous section, the method could be evaluated to examine if its performance was comparable to the model trained with the entire public dataset employed. In this section, we compared the performance of a well-trained YOLO v3 model to the difference between using the entire training dataset (without data augmentation techniques) and data partitioning (our approach). The same parameter choices were used in the approach’s training and testing. (Fig 14) depicts the confusion matrix for the normal approach. As (Fig 9(a) and Fig 14) are compared, it is evident that the YOLOv3 method (using the entire training dataset and without using data augmentation techniques) has higher prediction scores in the large two dataset classes, namely Neutrophil (segmented) and Myeloblast. While the approach detected one more class, Promyelocyte (bilobed), than ours when analyzing the testing dataset, the overall detection rate is lower.

**Fig 14.**
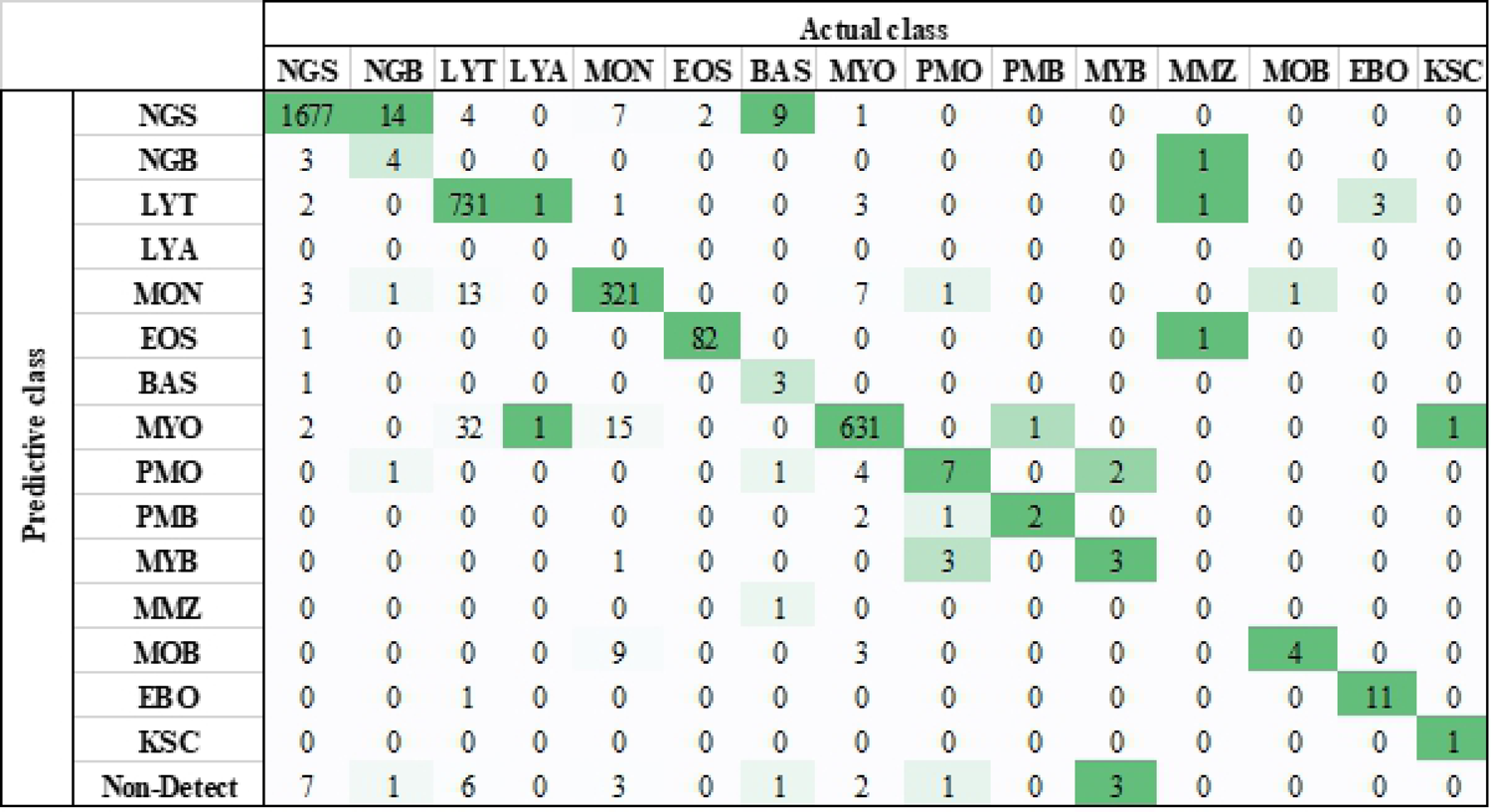
Confusion Matrix for YOLOv3 (Using Entire Training Dataset Without Applying Data Augmentation Techniques).

Table 2 further summarizes and contrasts the precision and sensitivity for multi-class classification of blood cell representations using a one-versus-rest method. In the precision contrast, the trained YOLOv3 model (using the entire testing dataset without using data augmentation techniques) has a higher precision score in five classes, while the trained model (our approach) has a higher precision score in seven classes. Both versions have higher sensitivity scores in six groups in the sensitivity comparison. As a result, our solution outperforms the competitive advantage in terms of precision and sensitivity for multi-class classification.

**Table 2.** Precision and Sensitivity Comparison in Multi-class Classification for the Two YOLOv3 Models. The better result is shown in bold.

| Classes | Precision |  | Sensitivity |  |
| --- | --- | --- | --- | --- |
|  | Entire Training Dataset +<br>Without Applying Data<br>Augmentation Techniques | Our<br>Approach | Entire Training Dataset +<br>Without Applying Data<br>Augmentation Techniques | Our<br>Approach |
| NGS | 0.978 | **0.988** | **0.989** | 0.974 |
| NGB | **0.5** | 0.273 | 0.19 | **0.429** |
| LYT | **0.985** | 0.984 | **0.929** | 0.926 |
| LYA | 0 | 0 | 0 | 0 |
| MON | **0.925** | 0.789 | 0.899 | **0.964** |
| EOS | 0.976 | **0.988** | **0.976** | 0.952 |
| BAS | 0.75 | **0.769** | 0.2 | **0.667** |
| MYO | 0.924 | **0.947** | **0.966** | 0.905 |
| PMO | 0.467 | **0.6** | 0.538 | **0.692** |
| PMB | **0.4** | 0 | **0.667** | 0 |
| MYB | 0.429 | **0.6** | 0.375 | 0.375 |
| MMZ | 0 | 0 | 0 | 0 |
| MOB | **0.25** | 0.222 | **0.8** | 0.4 |
| EBO | 0.917 | **0.929** | 0.786 | **0.929** |
| KSC | 1 | 1 | 0.5 | **1** |

Furthermore, we compared the overall performance of the two YOLOv3 models using micro-averaging, as shown in Table 3. According to the findings, the trained YOLOv3 model (using the entire training dataset without using data augmentation techniques) outperforms our approach in all performance metrics. At this point, our methodology has only used 33.51 % of the training dataset, and all of the performance are similar due to the slight calculation difference between the two models. The findings revealed that using data augmentation strategies in the training of a blood cell dataset would minimize the sample size from a broad dataset class whereas still being aided by the dataset necessity.

**Table 3.**
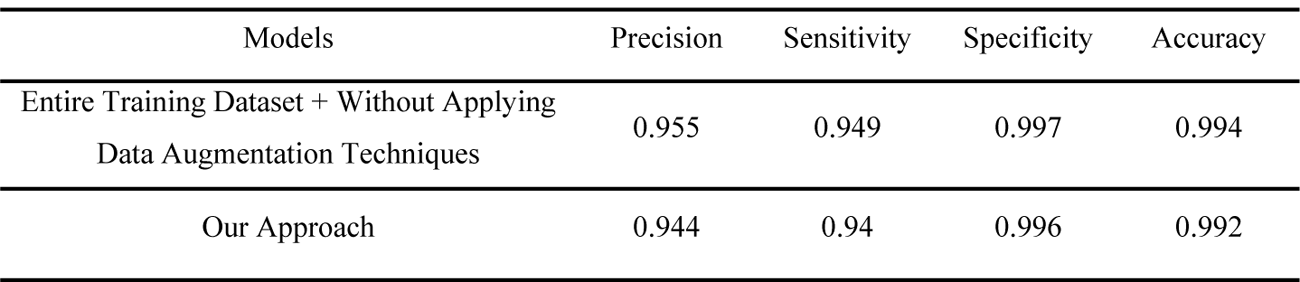
The Overall Performance Comparisons Between YOLOv3 (Using Entire Training Dataset Without Applying Data Augmentation Techniques) and YOLOv3 (Our Approach).

| Models | Precision | Sensitivity | Specificity | Accuracy |
| --- | --- | --- | --- | --- |
| Entire Training Dataset + Without Applying<br>Data Augmentation Techniques | 0.955 | 0.949 | 0.997 | 0.994 |
| Our Approach | 0.944 | 0.94 | 0.996 | 0.992 |

### ROC curve and the performance analysis on the varies of threshold level for the four YOLO approaches

For performance analysis using a ROC curve, four types of YOLO models are employed as classifiers to provide only a class decision, i.e., true class or false class on each instance. After applying these classifiers to a test dataset, it yields a confusion matrix for each threshold value which corresponds to one point in the ROC curve.

The classification results logically yield a numeric value of an instance probability with the predicted classes as shown in (Fig 10). The classifier produces the predicted classes if its output is higher than the predefined threshold values. From these results, we collected the overall values of four conditions such as TTP, TFP, TTN, and TFN scores of 15 classes from a confusion matrix. After that, we calculated the two important values (TPR and FPR) to plot a single point in the ROC curve. In this way, there are many different points in the ROC curve by varying the threshold values. Conceptually, we alter the threshold values from 0 (0%) to 1 (100%) to complete the ROC curve. Next, we performed the classification by using the testing dataset with the threshold values for each 0.05 (5%) increment. Consequently, we obtained the different 21 corresponding points in the ROC curve. The summarization of the classification results for four types of YOLO approach is shown in Table 4.

**Table 4.** The detailed calculations of TPR and FPR by using the four values (TTP, TFP, TTN, TFN) for each threshold values with the increment of 0.05 (5%) in four types of YOLO approaches.

| Threshold | YOLOv3 |  |  |  |  |  | YOLOv2 |  |  |  |  |  | YOLOv3-Tiny |  |  |  |  |  | YOLOv2-Tiny |  |  |  |  |  |
| --- | --- | --- | --- | --- | --- | --- | --- | --- | --- | --- | --- | --- | --- | --- | --- | --- | --- | --- | --- | --- | --- | --- | --- | --- |
|  | TTP | TFP | TTN | TFN | TPR | FPR | TTP | TFP | TTN | TFN | TPR | FPR | TTP | TFP | TTN | TFN | TPR | FPR | TTP | TFP | TTN | TFN | TPR | FPR |
| 0 | 3663 | 3663 | 0 | 0 | 1 | 1 | 3663 | 3663 | 0 | 0 | 1 | 1 | 3663 | 3663 | 0 | 0 | 1 | 1 | 3663 | 3663 | 0 | 0 | 1 | 1 |
| 0.05 | 3448 | 213 | 51069 | 215 | 0.94130 | 0.00415 | 3400 | 263 | 51019 | 263 | 0.92820 | 0.00513 | 3435 | 227 | 51055 | 228 | 0.93776 | 0.00443 | 3297 | 365 | 50917 | 366 | 0.90008 | 0.00712 |
| 0.1 | 3448 | 212 | 51070 | 215 | 0.94130 | 0.00413 | 3400 | 263 | 51019 | 263 | 0.92820 | 0.00513 | 3434 | 227 | 51055 | 229 | 0.93748 | 0.00443 | 3297 | 364 | 50918 | 366 | 0.90008 | 0.00710 |
| 0.15 | 3448 | 212 | 51070 | 215 | 0.94130 | 0.00413 | 3400 | 263 | 51019 | 263 | 0.92820 | 0.00513 | 3433 | 227 | 51055 | 230 | 0.93721 | 0.00443 | 3297 | 364 | 50918 | 366 | 0.90008 | 0.00710 |
| 0.2 | 3448 | 212 | 51070 | 215 | 0.94130 | 0.00413 | 3400 | 263 | 51019 | 263 | 0.92820 | 0.00513 | 3432 | 227 | 51055 | 231 | 0.93694 | 0.00443 | 3297 | 364 | 50918 | 366 | 0.90008 | 0.00710 |
| 0.25 | 3446 | 212 | 51070 | 217 | 0.94076 | 0.00413 | 3400 | 263 | 51019 | 263 | 0.92820 | 0.00513 | 3432 | 225 | 51057 | 231 | 0.93694 | 0.00439 | 3297 | 364 | 50918 | 366 | 0.90008 | 0.00710 |
| 0.3 | 3446 | 210 | 51072 | 217 | 0.94076 | 0.00410 | 3400 | 263 | 51019 | 263 | 0.92820 | 0.00513 | 3432 | 224 | 51058 | 231 | 0.93694 | 0.00437 | 3297 | 364 | 50918 | 366 | 0.90008 | 0.00710 |
| 0.35 | 3446 | 210 | 51072 | 217 | 0.94076 | 0.00410 | 3400 | 263 | 51019 | 263 | 0.92820 | 0.00513 | 3432 | 222 | 51060 | 231 | 0.93694 | 0.00433 | 3297 | 364 | 50918 | 366 | 0.90008 | 0.00710 |
| 0.4 | 3445 | 208 | 51074 | 218 | 0.94049 | 0.00406 | 3400 | 262 | 51020 | 263 | 0.92820 | 0.00511 | 3432 | 222 | 51060 | 231 | 0.93694 | 0.00433 | 3297 | 364 | 50918 | 366 | 0.90008 | 0.00710 |
| 0.45 | 3444 | 207 | 51075 | 219 | 0.94021 | 0.00404 | 3400 | 259 | 51023 | 263 | 0.92820 | 0.00505 | 3431 | 220 | 51062 | 232 | 0.93666 | 0.00429 | 3297 | 364 | 50918 | 366 | 0.90008 | 0.00710 |
| 0.5 | 3444 | 205 | 51077 | 219 | 0.94021 | 0.00400 | 3398 | 258 | 51024 | 265 | 0.92765 | 0.00503 | 3430 | 217 | 51065 | 233 | 0.93639 | 0.00423 | 3297 | 364 | 50918 | 366 | 0.90008 | 0.00710 |
| 0.55 | 3442 | 205 | 51077 | 221 | 0.93967 | 0.00400 | 3391 | 246 | 51036 | 272 | 0.92574 | 0.00480 | 3430 | 217 | 51065 | 233 | 0.93639 | 0.00423 | 3296 | 363 | 50919 | 367 | 0.89981 | 0.00708 |
| 0.6 | 3440 | 205 | 51077 | 223 | 0.93912 | 0.00400 | 3368 | 230 | 51052 | 295 | 0.91946 | 0.00449 | 3424 | 212 | 51070 | 239 | 0.93475 | 0.00413 | 3293 | 359 | 50923 | 370 | 0.89899 | 0.00700 |
| 0.65 | 3434 | 200 | 51082 | 229 | 0.93748 | 0.00390 | 3339 | 216 | 51066 | 324 | 0.91155 | 0.00421 | 3421 | 210 | 51072 | 242 | 0.93393 | 0.00410 | 3286 | 355 | 50927 | 377 | 0.89708 | 0.00692 |
| 0.7 | 3428 | 195 | 51087 | 235 | 0.93584 | 0.00380 | 3308 | 197 | 51085 | 355 | 0.90308 | 0.00384 | 3417 | 204 | 51078 | 246 | 0.93284 | 0.00398 | 3278 | 345 | 50937 | 385 | 0.89489 | 0.00673 |
| 0.75 | 3416 | 188 | 51094 | 247 | 0.93257 | 0.00367 | 3253 | 172 | 51110 | 410 | 0.88807 | 0.00335 | 3405 | 195 | 51087 | 258 | 0.92957 | 0.00380 | 3264 | 330 | 50952 | 399 | 0.89107 | 0.00644 |
| 0.8 | 3404 | 180 | 51102 | 259 | 0.92929 | 0.00351 | 3178 | 136 | 51146 | 485 | 0.86759 | 0.00265 | 3397 | 184 | 51098 | 266 | 0.92738 | 0.00359 | 3242 | 304 | 50978 | 421 | 0.88507 | 0.00593 |
| 0.85 | 3393 | 168 | 51114 | 270 | 0.92629 | 0.00328 | 3003 | 106 | 51176 | 660 | 0.81982 | 0.00207 | 3379 | 175 | 51107 | 284 | 0.92247 | 0.00341 | 3195 | 278 | 51004 | 468 | 0.87224 | 0.00542 |
| 0.9 | 3371 | 149 | 51133 | 292 | 0.92028 | 0.00291 | 2437 | 41 | 51241 | 1226 | 0.66530 | 0.00080 | 3357 | 158 | 51124 | 306 | 0.91646 | 0.00308 | 3140 | 234 | 51048 | 523 | 0.85722 | 0.00456 |
| 0.95 | 3309 | 122 | 51160 | 354 | 0.90336 | 0.00238 | 373 | 1 | 51281 | 3290 | 0.10183 | 0.00002 | 3306 | 132 | 51150 | 357 | 0.90254 | 0.00257 | 3031 | 156 | 51126 | 632 | 0.82746 | 0.00304 |
| 1 | 0 | 0 | 51282 | 3663 | 0 | 0 | 0 | 0 | 51282 | 3663 | 0 | 0 | 0 | 0 | 51282 | 3663 | 0 | 0 | 0 | 0 | 51282 | 3663 | 0 | 0 |

Considering the TPR and FPR results, we used the Jupyter Notebook open-source web application with a python environment to plot the ROC curve as shown in (Fig 15). In the ROC curve analysis, the AUC is an effective way to evaluate the performance of the trained model. The AUC value is always bounded between 0 and 1, where a perfectly inaccurate test represents a value of 0 and a perfectly accurate test represents a value of 1. In general, an AUC value can be defined as follows: under 0.5 is no realistic model, 0.5 is no discrimination model, 0.7 to 0.8 is considered acceptable model, 0.8 to 0.9 is considered excellent model, and more than 0.9 is considered outstanding model [38].

**Fig 15.**
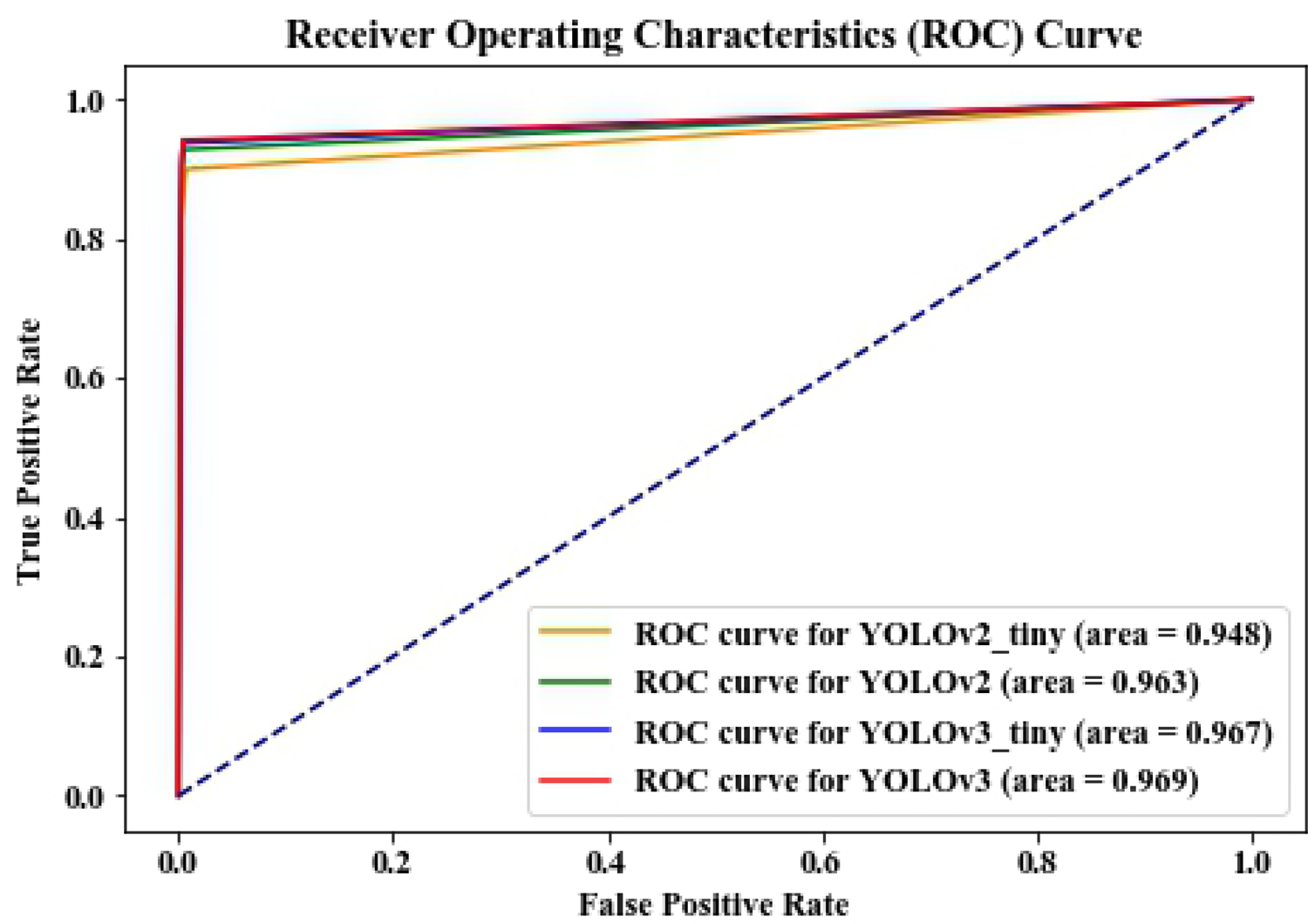
The ROC curve for four types of YOLO approach after evaluating testing dataset. The 21 data points of each YOLO approaches were received by adjusting the threshold values as shown in Table 4. The figure shows the corresponding ROC curve for four types of YOLO with the area under the curves (AUC) values.

According to (Fig 15), we conclude that the YOLOv3 model is a superior detection models because its AUC value is the highest one. The AUC values for the four YOLO models are 0.969, 0.967, 0.963, and 0.948 for YOLOv3, YOLOv3-Tiny, YOLOv2 and YOLOv2 respectively. Since all of the AUC values are higher than 0.9, we may consider the four models are an outstanding model for AML cell classifications.

We summarize our finding with the overall performance comparison between four types of YOLO approach as shown in Table 5. Although we varied the different threshold values for testing, we do not change NMS for all of the data analysis. In general, we may consider the four types of models have a good performance rating because of their performance scores. From Table 5, we found that the four performance values varied with the adjustment of threshold values. The result shows that the higher threshold values, the greater precision and specificity scores the models predicted.

**Table 5.** Overall performance comparison between the four types of YOLO approach for each 10% of threshold value increment.

| Performances | Model | Threshold |  |  |  |  |  |  |  |  |
| --- | --- | --- | --- | --- | --- | --- | --- | --- | --- | --- |
|  |  | 10% | 20% | 30% | 40% | 50% | 60% | 70% | 80% | 90% |
| Precision | YOLOv3 | 0.942 | 0.942 | 0.943 | 0.943 | 0.944 | 0.944 | 0.946 | 0.950 | 0.958 |
|  | YOLOv2 | 0.928 | 0.928 | 0.928 | 0.928 | 0.929 | 0.936 | 0.944 | 0.959 | 0.983 |
|  | YOLOv3-Tiny | 0.938 | 0.938 | 0.939 | 0.939 | 0.940 | 0.942 | 0.944 | 0.949 | 0.955 |
|  | YOLOv2-Tiny | 0.901 | 0.901 | 0.901 | 0.901 | 0.901 | 0.902 | 0.905 | 0.914 | 0.931 |
| Sensitivity | YOLOv3 | 0.941 | 0.941 | 0.941 | 0.940 | 0.940 | 0.939 | 0.936 | 0.929 | 0.920 |
|  | YOLOv2 | 0.928 | 0.928 | 0.928 | 0.928 | 0.928 | 0.919 | 0.903 | 0.868 | 0.665 |
|  | YOLOv3-Tiny | 0.937 | 0.937 | 0.937 | 0.937 | 0.936 | 0.935 | 0.933 | 0.927 | 0.916 |
|  | YOLOv2-Tiny | 0.900 | 0.900 | 0.900 | 0.900 | 0.900 | 0.899 | 0.895 | 0.885 | 0.857 |
| Specificity | YOLOv3 | 0.996 | 0.996 | 0.996 | 0.996 | 0.996 | 0.996 | 0.996 | 0.996 | 0.997 |
|  | YOLOv2 | 0.995 | 0.995 | 0.995 | 0.995 | 0.995 | 0.996 | 0.996 | 0.997 | 0.999 |
|  | YOLOv3-Tiny | 0.996 | 0.996 | 0.996 | 0.996 | 0.996 | 0.996 | 0.996 | 0.996 | 0.997 |
|  | YOLOv2-Tiny | 0.993 | 0.993 | 0.993 | 0.993 | 0.993 | 0.993 | 0.993 | 0.994 | 0.995 |
| Accuracy | YOLOv3 | 0.992 | 0.992 | 0.992 | 0.992 | 0.992 | 0.992 | 0.992 | 0.992 | 0.992 |
|  | YOLOv2 | 0.990 | 0.990 | 0.990 | 0.990 | 0.990 | 0.990 | 0.990 | 0.989 | 0.977 |
|  | YOLOv3-Tiny | 0.992 | 0.992 | 0.992 | 0.992 | 0.992 | 0.992 | 0.992 | 0.992 | 0.992 |
|  | YOLOv2-Tiny | 0.987 | 0.987 | 0.987 | 0.987 | 0.987 | 0.987 | 0.987 | 0.987 | 0.986 |

## Conclusions

We introduce and evaluate the four types of the well-known object detection strategy, YOLO algorithm to detect the 15-class of WBC cell images. Data augmentation techniques are also applied to enhance and adjust the training images in the dataset. These include the basic image processing procedures of rotation, contrast, noise and blur. After performing the four image processing tasks, one image can be augmented into up to 108 images. The training dataset with 608×608 pixels image was further preparation for the training dataset by these augmented images. The four models of YOLO approach (YOLOv3, YOLOv3-Tiny, YOLOv2 and YOLOv2-Tiny) are used for the classification of Acute Myeloid Leukaemia (AML) Blood Cell. YOLOv3 exhibits a more reliable performance than the other three models. YOLOv3 demonstrates 94% in overall precision and sensitivity and 99% in overall specificity and accuracy. Furthermore, even though we used 33.51 percent of the training data in model training, our suggested YOLOv3 solution outperforms the other YOLOv3 approach (using the whole training dataset and without using data augmentation techniques). We also propose the performance evaluation procedure using the ROC curve. This procedure can be used to examine the performance of our approaches with quantitative AUC values. The AUC values for the four YOLO models are 0.969, 0.967, 0.963, and 0.948 for YOLOv3, YOLOv3-Tiny, YOLOv2 and YOLOv2-Tiny respectively. Since the obtained AUC scores are higher than 0.9, we can consider the four models are outstanding models for AML cell classifications. With above performance, our approach using deep learning can potentially make more reliable and faster clinical diagnoses in the future.

## Acknowledgements

This work was supported by Thailand Science Research and Innovation Fund and King Mongkut’s Institute of Technology Ladkrabang (KMITL), Thailand. We acknowledge with thanks to The Cancer Imaging Archive (TCIA) for their publicly available dataset.

## Author Contributions

**Conceptualization:** Kaung Myat Naing, Veerayuth Kittichai, Siridech Boonsang.

**Data collection:** Veerayuth Kittichai, Kaung Myat Naing.

**Funding acquisition:** Siridech Boonsang.

**Investigation:** Kaung Myat Naing, Veerayuth Kittichai.

**Methodology:** Kaung Myat Naing.

**Project administration:** Veerayuth Kittichai.

**Software:** Teerawat Tongloy, Kaung Myat Naing.

**Supervision:** Veerayuth Kittichai, Santhad Chuwongin, Siridech Boonsang.

**Writing – original draft:** Kaung Myat Naing, Veerayuth Kittichai.

**Writing – review & editing:** Veerayuth Kittichai, Santhad Chuwongin, Siridech Boonsang.

